# Fast, live-cell imaging of 15 intracellular compartments by deep learning segmentation of super-resolution data

**DOI:** 10.1101/2021.12.13.472520

**Authors:** Karl Zhanghao, Meiqi Li, Xingye Chen, Wenhui Liu, Yiming Wang, Zihan Wu, Chunyan Shan, Jiamin Wu, Yan Zhang, Peng Xi, Dayong Jin

## Abstract

The number of colors that can be used in fluorescence microscopy to image the live-cell anatomy and organelles’ interactions is far less than the number of intracellular organelles and compartments. Here, we report that deep convolutional neuronal networks can predict 15 subcellular structures from super-resolution spinning-disk microscopy images using only one dye, one laser excitation, and two detection channels. Comparing to the colocalization images, this method achieves pixel accuracies of over 91.7%, which not only bypasses the fundamental limitation of multi-color imaging but also accelerates the imaging speed by more than one order of magnitude.

## Maintext

Until now, up to six colors excited by multiple lasers in fluorescence imaging can be used to simultaneously observe the interaction and coordination between organelles^1-4^. Further increase in color will be intrinsically limited by the dyes’ cross-talks in the spectrum domain, requiring special design of probes^5^. Besides, the labeling procedure becomes tedious and their labeling efficiency will drop exponentially when increasing the types of fluorophores. The long duration, as a result of the multiple excitations and image acquisition steps, is essentially required in multi-color fluorescence imaging, and consequently the phototoxicity further becomes a concern in live-cell imaging. Deep learning has been recently introduced to in-silico prediction of multi-color images from transmitted bright field images^6-9^. Nevertheless, they are all spatially diffraction-limited, suffering from low resolution and low contrast, therefore the prediction accuracy in recognizing the intracellular organelles is yet to be satisfactory^6^.

Super-resolution microscopy, by taking advantage of their high resolution and contrast, has enabled the live-cell imaging of subcellular structures and dynamics^10-13^. Here we explore the segmentations of super-resolution images obtained from a commercial super-resolution spinning disk microscope (Fig. S1)^14^. To simplify both the labeling process and image collections, we universally stain all the subcellular lipid membranes with a lipid dye Nile Red. We further employ a two-color detection to discriminate vesicle organelles with similar shapes and sizes, as the emission spectrum of the dye responds to the lipid polarity of membranes^14^. We demonstrate such a simple preparation of cell staining and rapid acquisition of spatial and spectral imaging data can empower deep convolutional neural network (DCNN) to predict intracellular anatomy images with high accuracy and throughput, providing a new paradigm for multiplexing imaging inside the living cells.

Learning from the dual-color super-resolution images, our DCNN developed with multiple sub-models can predict 15 intracellular compartments (Fig. 1, supplementary Movie 1). Namely, LD-net is responsible for recognizing lipid droplets, GOLGI-net for Golgi apparatus, MITO-net for mitochondria, PERO-net for peroxisome, EE-net for early endosome, LE-net for late endosome, LYSO-net for lysosome. ER-net is used to recognize the three structures of endoplasmic reticulum (ER), nuclear membrane, and nuclear reticulum, PM-net is used to recognize plasma membrane and filopodia, and VOLUME-net is to recognize other three structures of nucleus, cytosol, and extracellular space (ECS).

**Fig. 1.**
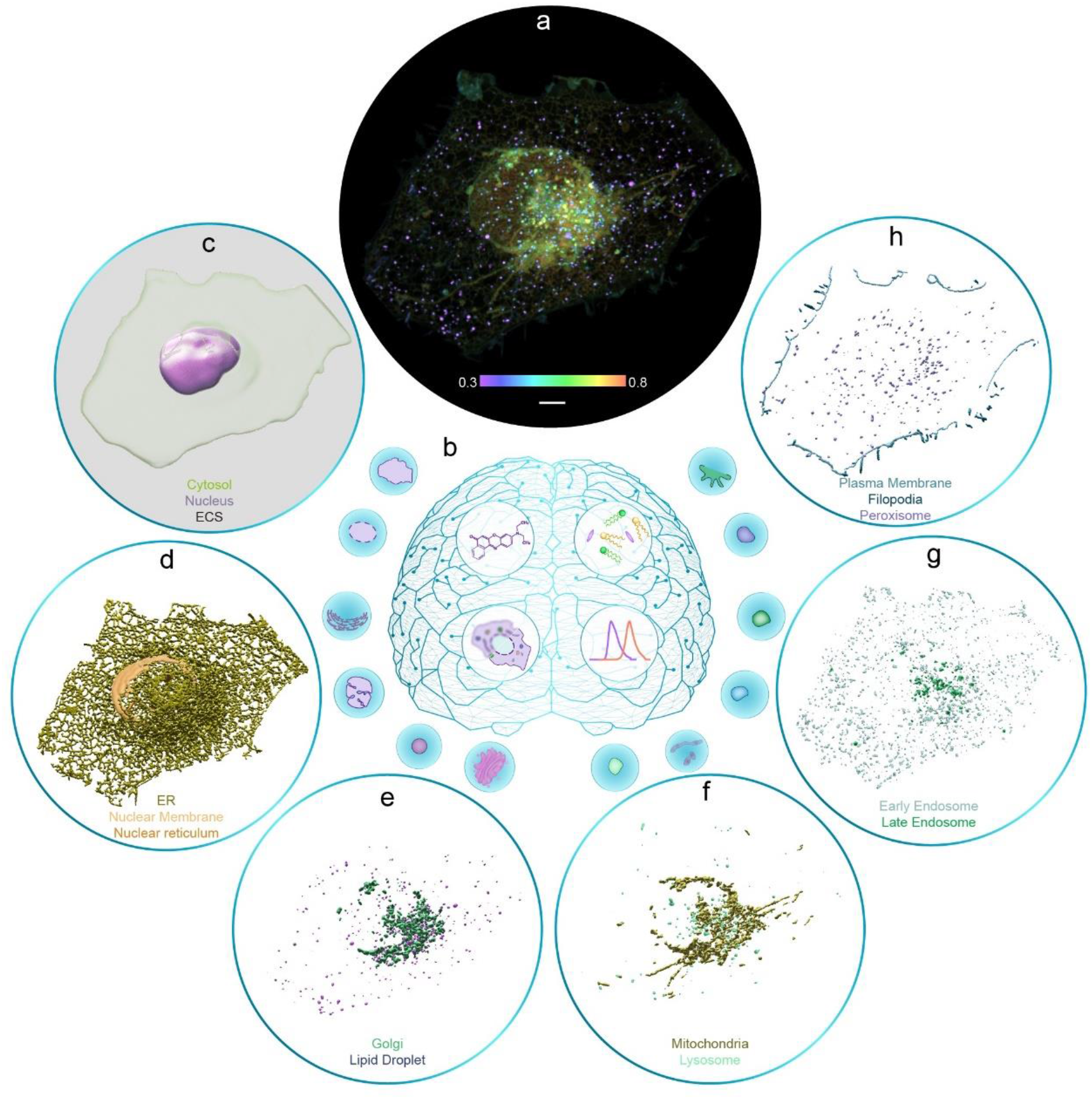
Imaging cell anatomy via super-resolution microscopy and deep learning. **(a)** Super-resolution image of a U2-OS cell with its membranous structures stained by Nile Red. The pseudo color represents the spectral ratio between the yellow channel (Em: 580-653 nm) and the red channel (Em: 665-705 nm) under single laser excitation at 488 nm. **(b)** From the super-resolution intensity image and the spectral ratio image, the deep convolutional neural networks predict binary masks of 15 subcellular structures. **(c)** The masks of cytosol, nucleus, and extracellular space segmented by VOLUME-net. **(d)** The masks of ER, nuclear reticulum, and nuclear membrane segmented by ER-net. **(e)** Golgi mask segmented by GOLGI-net and lipid droplet mask segmented by LD-net. **(f)** Mitochondria mask segmented by MITO-net and lysosome mask segmented by LYSO-net. **(g)** Early endosome mask segmented by EE-net and late endosome mask segmented by LE-net. **(h)** Peroxisome mask segmented by PERO-net and the masks of plasma membrane and filopodia segmented by PM-net. Scale bar: 5 μm.

We segment each structure with a binary mask, which can further multiply the Nile Red image to obtain a corresponding intensity image (Fig. 2a, b). Compared with the prediction of multiple intensity levels, binary prediction simplifies the problem with true/false decisions and avoids the generation of fake signals. The binary masks are also sufficient for further quantitative analysis, such as organelles’ number, volume, and contact frequency, which can provide a full picture of the cell anatomy.

**Fig. 2.**
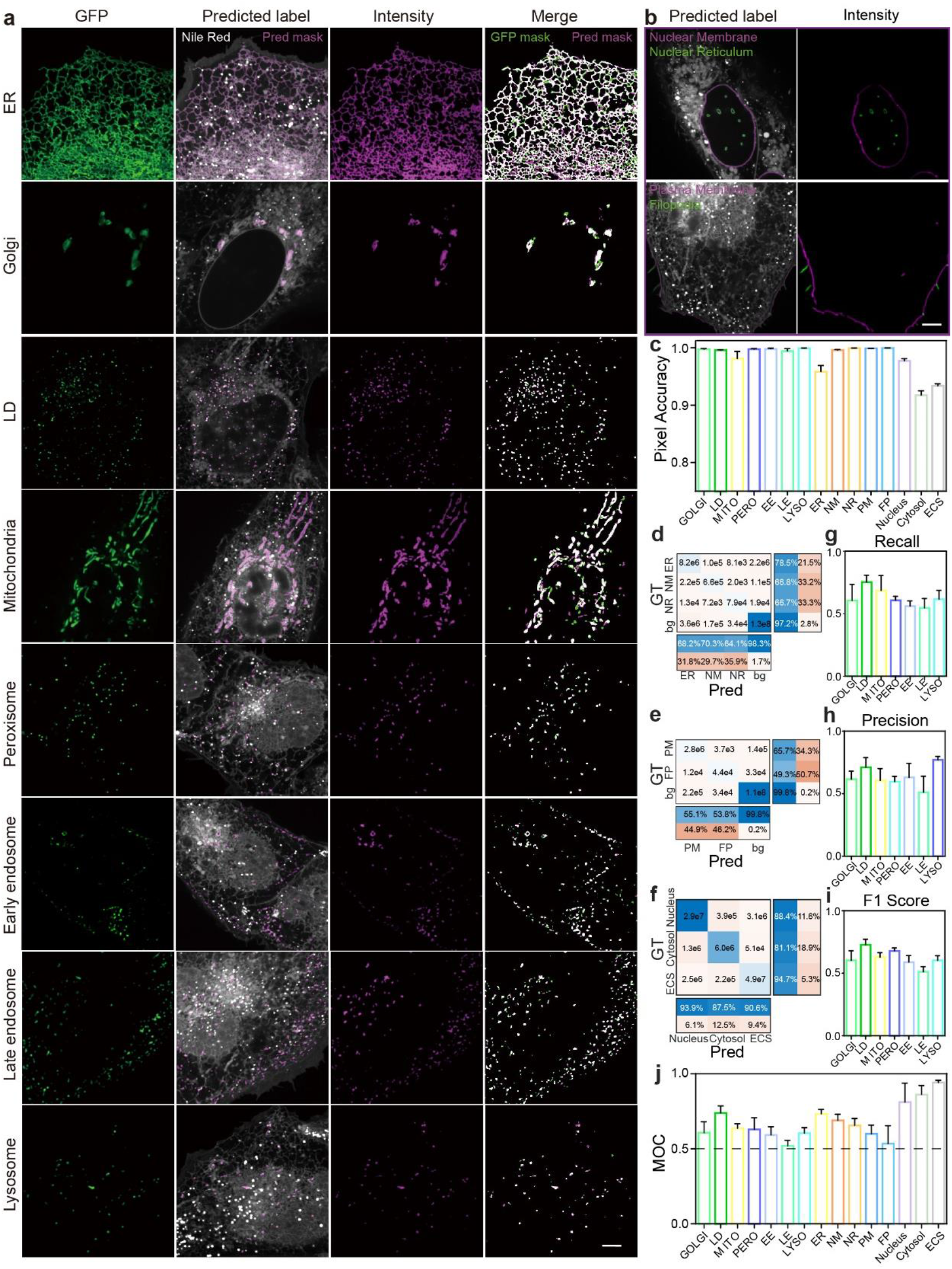
Comparison between the network predicted images and colocalization images. **(a)** The first column is colocalization images; the second column is Nile Red fluorescence images with magenta masks predicted by the networks; the third column is intensity images by multiplying the fluorescence image and the predicted masks; the fourth column is merged images of predicted masks (in magenta) and colocalization masks (in green, FG-net prediction of colocalization intensity images). In the merged images, magenta pixels indicate wrong prediction (false positive) and green pixels indicate missing prediction (false negative). **(b)** The predicted masks of nuclear membrane, nuclear reticulum, plasma membrane, and filopodia on the fluorescence images and the masked intensity images. **(c)** Pixel accuracies of 15 structures. **(d-f)** Confusion matrixes of multi-classification networks including ER-net, PM-net, and VOLUME-net. **(g-i)** Recall, precision, and f1 score of binary segmentation networks. **(j)** The Manders’ overlapping coefficient of 15 structures between predicted masks and ground-truth masks. Scale bar: 5μm.

To train the DCNN networks, the masks for nuclear membrane, nuclear reticulum, plasma membrane, filopodia, nucleus, cytosol, and ECS, are manually annotated on the super-resolution images. For other compartments, we acquire a green-channel colocalization image with organelle-specific probes. Since Nile Red has a broad emission band, we also use linear unmixing and the output of the pre-trained LD-net to exclude the Nile Red fluorescence in the colocalization channel (online Methods). An FG-net (Fig. S2b) has been trained to segment the colocalization image separating the foreground (in-focus) area from the background (out-focus or dark) area. Unlike conventional foreground extraction methods based on thresholding, where manual adjustment of parameters is needed for different structures, our deep learning approach not only bypasses the image preprocessing and parameter adjustments but also shows superior performance in predicting masks of various organelles (Fig. S3).

The super-resolution images and ground truth binary masks are used to train networks based on the attention U-Net architecture^15, 16^ (Fig. S2a). The input images are resized and cropped into 3D patches matching the size of a cell. Golgi-net, LD-net, MITO-net, PERO-net, EE-net, LE-net, and LYSO-net are binary segmentation networks optimized with sigmoid cross entropy loss, while ER-net, PM-net, and VOLUME-net are multi-class segmentation networks optimized with softmax cross entropy loss. Since the abundance of different structures varies, we adjust the class weights for different networks. Details about the training datasets are included in Supplementary Table 1.

As shown in Fig. 2a, b, the comparison between the typical network predicted images and the colocalization images shows high accuracies for these intracellular compartment structures. Fig. 2c further quantitatively reports the overall pixelwise accuracies of higher than 91.7%. Moreover, confusion matrixes are used to evaluate multi-class segmentation networks of ER-net, PM-net, and VOLUME-net (Fig. 2d-f), which reveals the large portion of true positive and true negative pixels. For binary segmentation networks, the recall, precision, and F1 score (Fig. 2g-i) are used to indicate their accurate prediction. From each structure, by calculating the Mander’s overlapping coefficient (MOC)^17^ between the predicted mask and the ground truth mask, masks of ER, lipid droplet, cytosol, nucleus, and ECS display strong colocalizations with ground truth (MOC>0.7); masks of all other structures show good colocalizations (MOC>0.5) (Fig. 2h).

To explore the opportunity offered by the ratio images in segmentation accuracies, we found that the ratio image was critical for achieving the accurate segmentations of vesicle organelles with similar size and shape. Fig. S4b shows that the prediction accuracy of early endosome is much higher with the additional ratio image (F1score=0.59 vs 0.43). But for other structures, such as mitochondria, the additional ratio images did not add much value to the prediction accuracy, compared with the result by only using the intensity image (F1score=0.63 vs 0.62, Fig. S4a). Therefore, the segmentation accuracies rely on both super-resolution morphologies and organelle-specific spectrum ratios. In comparison, the DCNN network with bright field images predicted the Golgi image with a low Pearson’s correlation coefficient (PCC<0.2)^6^, while our segmentation of the Golgi mask delivers a much better result of MOC=0.61.

Here we demonstrate the multiplexed and highly accurate deep learning segmentation on a widely accessible LiveSR microscope. With this technique we further imaged the cell anamoty at different mitosis stages (Fig. S5, Supplementary Movie 2-7). Imaging subcellular structures using Nile Red staining has also been demonstrated with other super-resolution microscopy, including Structured Illumination Microscopy (SIM) ^14^, STimulated Emission Depletion (STED) ^18^, or Single Molecule Localization Microscopy (SMLM) ^19^, which may also benefit the DCNN based segmentation approach. Our technique can be easily expanded in association with other fluorescent probes, to further study the subcellular interactions among organelles, cytoskeleton, and proteins.

## Methods

### Sample Preparation

Human osteosarcoma U2-OS cell lines (HTB-96, ATCC, USA) were cultured in Dulbecco’s Modified Eagle’s medium (DMEM, GIBCO, USA) containing 10% heat-inactivated fetal bovine serum (FBS, GIBCO, USA) and 100 U/ml penicillin and 100 μg/ml streptomycin solution (PS, GIBCO, USA) at 37°C in an incubator with 95% humidity and 5% CO_2_. For the living cell imaging, the cells were plated at the desired density on the μ-Slide 8 Well (80827, ibidi, USA) and 1 μg/ml Nile Red (N1142, Invitrogen, USA) was added into the culture medium 1 h before imaging and was present during imaging. For colocalization experiments, the cells were transfected 16 h before imaging with the plasmids of early endosome-GFP (Rab5a, C10586, BacMam 2.0, CellLight, USA), late endosome-GFP (Rab7a, C10588), ER-GFP(ER signal sequence of calreticulin and KDEL, C10590), Golgi-GFP (Golgi-resident enzyme N-acetylgalactosaminyltransferase 2, C10592), Mitochondria-GFP (leader sequence of E1 alpha pyruvate dehydrogenase, C10600) and incubated overnight. 1 μg/ml Nile Red was added into the culture medium 1 h before imaging. The colocalization of lysosome is obtained by Lysoview™ 488 (70067-T, Biotium) 30min before imaging and without washing during imaging.

### System setup

The data were acquired from a commercial system based on the inverted fluorescence microscope (TI-E, Nikon) equipped with a TIRF objective (CFI Apochromat TIRF ×100 oil, NA 1.49, Nikon) and a spinning disk confocal system (CSUW1, Yokogawa). The super-resolution imaging module (Live SR, Gataca) could double the imaging resolution. One laser (iLAS 3, 488 nm-150mW) is used for excitation. Emission fluorescence was acquired by sCMOS camera (Prime 95B, Photometrics) after a multi-splitter module (CAIRN, MultiSplit V2) with three detection channels (green: S525/50m; yellow: S685/40m; red: S617/73m; Chroma). The acquisition process is performed on the Metamorph software. The resolution of the system is analyzed by PSFj software (Fig. S1).

### Training data preparation

Multi-channel images are registered with chromatic aberration correction algorithm^20^. The emission ratio was calculated by dividing the red channel by the yellow channel. The fluorescence image is the average image between the red channel and yellow channel. The whole imaging FoV is 1200*1200*Nz px^3^ (Nz ranges from 12 to 30), which is resized into 802*802*24 px^3^ and is further cropped into random 256*256*24 px^3^ patches.

For the structures of nuclear membrane, nuclear reticulum, plasma membrane, filopodia, nucleus, cytosol, and ECS, the ground truth masks are manually annotated. For other structures, the green-channel colocalization images are acquired. In green-channels images, Nile Red shows strong fluorescence in LD and weak fluorescence in other organelles. To exclude the week Nile Red fluorescence in other organelles, the green-channel images are subtracted by a portion of the yellow-channel images (ChG-r*ChY, r=0.2). To exclude the strong Nile Red fluorescence in LD, the pixels within the LD mask, which is predicted by the pre-trained LD network, are set to zero intensity. The ground truth masks are segmented from the colocalization image by a pre-trained FG-net (Fig. S2b), which is trained with 40 manually annotated colocalization images of various organelles.

### Model architecture and training

Our DCNN networks are based on the attention U-Net architecture (Fig. S2a) and are implemented in Python using the PyTorch package. The FG-net takes 2D intensity images as input (1@256*256), and the subcellular segmentation networks take 3D intensity and ratio images as input (2@256*256*24). Both binary segmentation networks and multi-class segmentation networks are optimized by Adam optimizer with cross entropy loss functions. The starting learning rate of 0.0001, which is reduced on plateau with the factor of 0.3 and patience of 5. The class weights of different organelles are adjusted according to the abundance of corresponding organelles. For example, a cell consists of more pixels labeled as mitochondria than as peroxisome, so that the weight of mitochondria is 2 and the weight of peroxisome is 5 (Supplementary Table 1). The number of training patches varies from 160∼204 for different structures, which are trained on the RTX 2080 Ti GPU within 12 h.

### Image analysis and display

The evaluation of predicted masks is performed by custom-written Matlab. The predicted masks for each patch are merged and resized back, and the metrics between the ground truth and prediction are calculated as follows:

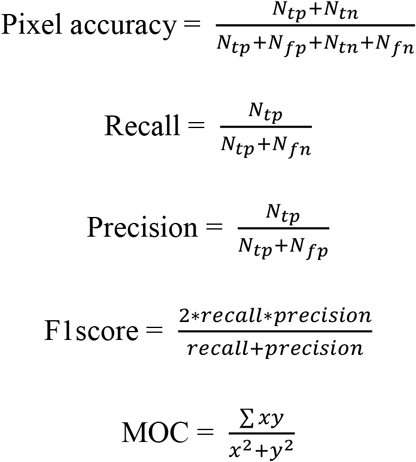

where *N*_*tp*,_*N*_*fp*,_*N*_*tn*,_*N*_*fn*,_are the number of pixels for true-positive, false-positive, true-negative, and false-negative; *x,y* are pixel values of the predicted mask and the ground truth mask. For the confusion matrixes, the value in the *i*th row, *j*th column (*N*_*ij*_) is the number of pixels with label *i* in the ground truth image and label *j* in the predicted image. The 3D intensity images are generated in Imaris (https://imaris.oxinst.com/) by volumetric intensity rendering and the 3D volume of binary masks is generated by surface rendering.

## Supporting information

Visualization of 15 subcellular structures within a single cell

